# Temporal information of subsecond sensory stimuli in primary visual cortex is encoded via high dimensional population vectors

**DOI:** 10.1101/2024.01.05.574126

**Authors:** Sam Post, William Mol, Noorhan Rahmatullah, Anubhuti Goel

## Abstract

Whether in music, language, baking, or memory, our experience of the world is fundamentally linked to time. However, it is unclear how temporal information is encoded, particularly in the range of milliseconds to seconds. Temporal processing at this scale is critical to prediction and survival, such as in a prey anticipating not only where a charging predator will go but also *when* the predator will arrive at that location. Several models of timing have been proposed that suggest that either time is encoded intrinsically in the dynamics of a network or that time is encoded by mechanisms that are explicitly dedicated to temporal processing. To determine how temporal information is encoded, we recorded neural activity in primary visual cortex (V1) as mice (male and female) performed a goal directed sensory discrimination task, in which patterns of subsecond stimuli differed only in their temporal profiles. We found that temporal information was encoded in the changing population vector of the network and that the space between these vectors was maximized in learned sessions. Our results suggest that temporal information in the subsecond range is encoded intrinsically and does not rely upon specialized timing mechanisms.

**SIGNIFICANCE STATEMENT:** Our experience of the world is fundamentally linked to time, but it is unclear how temporal information is encoded, particularly in the range of milliseconds to seconds. Using a sensory discrimination task in which patterns of subsecond stimuli differed in their temporal profiles, we found that primary visual cortex encodes temporal information via the changing population vector of the network. As temporal processing via population encoding has been shown to rely on inhibitory activity in computational models, our results may provide insight into temporal processing deficits in disorders such as autism spectrum disorder in which there is inhibitory-excitatory imbalance. Furthermore, our results may underlie processing of higher-order sensory stimuli, such as language, that are characterized by complex temporal sequences.

## INTRODUCTION

Our experience of the world is fundamentally linked to time. We rely upon its even structure and passage and are as a result, able to make predictions about the future. We anticipate winter following autumn, and we know that the sun will set and then rise again. When we are driving, we expect a red light to follow a yellow, and a green light to follow a red. The structure of these events is sequential, which is not inherently connected with time, but within each sequence there is a temporal dimension. For instance, we decide to press the brake or the gas pedal based on our estimation of the duration of the yellow light. And we would be quite concerned if one day the sun rose ten minutes after setting, or perhaps if night spontaneously stretched out for several years.

Neuroscience has made great progress in elucidating how sensory and motor content are encoded, whether in the present, such as during stimulus discrimination, or in the past, such as in memory encoding. Time’s role in these encoding schemes has been largely overlooked however, which may simply be the result of its ubiquity. There is no sensory organ that measures time, though in each sensory modality time is present. This realization then begs the question of how time is encoded: might it be encoded intrinsically within each sensory modality, or is it encoded by higher order mechanisms specifically dedicated to it?

Increasingly, evidence points to a variety of mechanisms, and these largely depend upon the scale of an interval. On the order of days, transcriptional feedback loops in the suprachiasmatic nucleus are responsible (Mauk & Buonomano, 2004). On the order of minutes, corticostriatal loops mediated by dopaminergic activity are the likely mechanism (Mauk & Buonomano, 2004). However, on the order of seconds and milliseconds, the mechanisms of temporal encoding remain unclear and widely debated.

The importance of timing at this scale is acutely linked to prediction and survival. A boxer anticipates at what moment to slip their opponent’s punch, and a prey watching a charging predator must predict not only where a predator will go, but also at what moment the predator arrives at that location. However, temporal encoding at this scale is not simply limited to interval timing (i.e. the duration of a stimulus, or the duration between two stimuli) like in these examples, but undergirds an array of simple to complex phenomena. Indeed, temporality is endemic to highly complex stimuli such as music, Morse code, and language, in which meaning is intrinsically derived from temporal structure.

Several models of timing at this scale have been proposed that largely lie on a dedicacated to intrinsic axis (**Fig. 1**), but it remains to be determined which accounts best for temporal encoding of sensory stimuli. Here, we investigate how subsecond temporal information is encoded in V1 in a goal directed sensory discrimination task, in which temporal information exclusively differentiates stimuli. We previously showed that mice become experts at the task and that changes in V1 dynamics accompany expert performance in the learned session (Post et al., 2023). In this paper, we show the evolution of neural dynamics through learning and test whether dedicated or intrinsic mechanisms are employed in temporal encoding of sensory stimuli. We find that temporal information is encoded in the changing population vector, i.e. trajectory, of the network through high dimensional space. This finding evinces a prominent intrinsic model of timing, the state dependent network model. Additonally, we find that neural activity which may be representative of dedicated models of timing, namely ramping and oscillatory models, is no more representative of temporal information than non-specialized activity and is in fact an aspect of the changing population vector in state space. Our results add to a growing body of literature which suggests that temporal information is intrinsically encoded in the processing of sensory stimuli.

**FIGURE 1:**
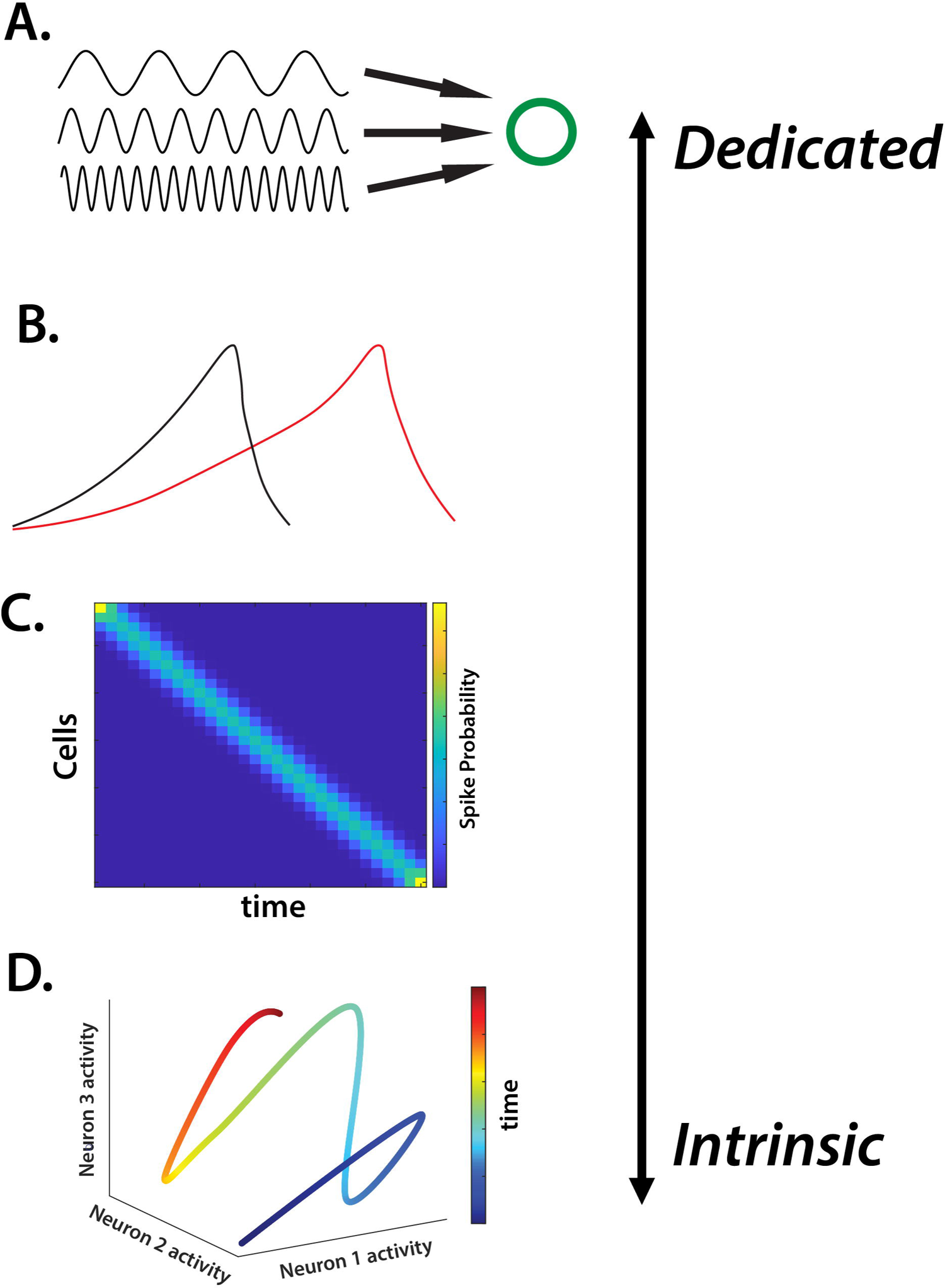
Axis of dedicated to intrinsic models of timing models. **A.** Oscillatory models: intrinsic oscillators project to a downstream readout unit or units. Early models proposed that a pacemaker, much like a metronome, output periodic pulses which were then counted by a downstream accumulator (Creelman, 1962; Gibbon, 1977); recent oscillatory models propose that a network of cortical oscillators with differing periodicities project to striatal medium spiny neurons that act as coincidence detectors and decode the oscillatory output (Buhusi & Meck, 2005; Matell & Meck, 2004; Merchant et al., 2013, 2015; van Rijn et al., 2014). **B.** Ramping models: temporal information is encoded by the firing rate of a given neuron and were motivated by work in decision making in non-human primates (de Lafuente et al., 2022). Ramping activity has been found in sensory areas like primary visual cortex (V1) (Chubykin et al., 2013; Monk et al., 2020; Shuler & Bear, 2006). **C.** Synfire chain models: activity is sparsely tiled over a population. **D.** State dependent network models: temporal information is encoded in the changing population vector, i.e. trajectory, of a network through high dimensional state space. A simple 3 unit network illustrates how the state of the network changes in 3 dimensional space over time. Experimental and computational evidence increasingly points to the state dependent model as the candidate mechanism of temporal encoding on the order of milliseconds to seconds (Buonomano, 2000; Goel & Buonomano, 2016; Karmarkar & Buonomano, 2007; Post et al., 2023; Seay et al., 2020; Zhou et al., 2022).

## MATERIALS AND METHODS

### Experimental Animals

All experiments followed the U.S. National Institutes of Health guidelines for animal research, under animal use protocols approved by the Chancellor’s Animal Research Committee and Office for Animal Research Oversight at the University of California, Riverside (ARC #2022-0022). We used male and female FVB.129P2 WT mice (JAX line 004828). All mice were housed in a vivarium with a 12/12 h light/dark cycle and experiments were performed during the light cycle. The FVB background was chosen because of its robust breeding. 4 males and 1 female were used.

### Go/No-go temporal pattern sensory discrimination (TPSD) task for head restrained mice

Awake, head-restrained young adult mice (2-4 months) were allowed to run on an air-suspended polystyrene ball while performing the task in our custom built rig (**Fig. 2B**). Prior to performing the task, the animals were subjected to handling, habituation, and pretrial phases. After recovery from headbar/cranial window surgery, mice were handled gently for 5 min every day, until they were comfortable with the experimenter and would willingly transfer from one hand to the other to eat sunflower seeds. This was followed by water deprivation (giving mice a rationed supply of water once per day) and habituation to the behavior rig. During habituation, mice were head-restrained and acclimated to the enclosed sound-proof chamber and allowed to run freely on the 8 cm polystyrene ball. Eventually, mice were introduced to the lickport that dispensed water (3-4 µL) and recorded licking (custom-built at the UCLA electronics shop), followed by the audio-visual stimuli. This was repeated for 10 min per session for 3 days. Starting water deprivation prior to pretrials motivated the mice to lick (Guo et al., 2014). After habituation and ∼15% weight loss, mice started the pretrial phase of the training. During pretrials, mice were shown the Pref stimulus only with no punishment time associated with incorrect responses. This was done in order to teach the mice the task structure and encourage the mice to lick and to remain motivated. The first day consisted of 150 trials and subsequent days of 250. The reward, as in the TPSD main task, was dispensed at 1.2 s and remained available to the mice until 2 s, at which time it was sucked away by a vacuum. The mice were required to learn to associate a water reward soon after the stimulus was presented and that there was no water reward in the inter-trial interval (4 s period between trials). Initially during pre-trials, the experimenter pipetted small drops of water onto to the lickport to coax the mice to lick. Once the mice learned this and licked with 80% efficiency, they were advanced to the go/no-go task.

The TPSD task is a go/no-go task composed of two sequences of synchronous audio-visual stimuli (**Fig. 2A**). Visual stimuli are 90° drifting sinusoidal gratings and are accompanied by a synchronous 5 kHz tone at 80 dB. Within each sequence, four stimuli are presented that differ only in temporality. Our preferred sequence is composed of 4 stimuli of 200 ms; our nonpreferred sequence is composed of 4 stimuli of 900 ms. Each set of the sequences is separated by a 200 ms period of silence accompanied by a grey screen. A water reward is dispensed at 1.2 s and remains available until 2 s, at which time it is sucked away by a vacuum. A custom built lickport (UCLA engineering) dispensed water, vacuumed it, and recorded licking via breaks in an infrared (IR) beam. Breaks were recorded at 250 Hz. The window in which mice’s licking count toward a response is 1 to 2 s from stimulus onset in both conditions. A time out period (6.5 to 8 s), in which the monitor shows a black screen and there is silence, is instituted if the mouse incorrectly responds. The first session was composed of 250 trials, and subsequent days of 350. Depending on the stimulus presented, the animal’s behavioral response was characterized as “Hit”, “Miss”, “Correct Rejection” (CR) or “False Alarm” (FA) (**Fig. 2A**). An incorrect response resulted in the time-out period.

To expedite learning, we set the ratio of preferred to nonpreferred stimuli to 70:30 as we found that mice are more prone to licking (providing a ‘yes’ response) than to inhibiting licking (providing a ‘no’ response). We additionally instituted an individualized lick rate threshold to encourage learning as we found that lick rates differed significantly from mouse to mouse. Licking thresholds were calculated from lick rates for mice and shows no significant correlation between licking thresholds and learning rates (Pearson’s *r*, r = .4684, p = -.3012). This indicates that the individualized lick rate threshold was used as a learning aid to complete the task and did not affect their learning rates or their reliance on the stimulus for task completion. To confirm that mice learned rather than took advantage of the biased 70:30 preferred to nonpreferred trial ratio, we tested mice for 2 additional sessions using a 60:40 ratio of preferred to nonpreferred. We retain a greater number of preferred stimuli as the total time mice encounter preferred stimuli is less than that of encountering nonpreferred stimuli within a 60:40 trial session (294 s vs 588 s respectively). Following, mice performed a control task, during which visual and auditory stimuli were not presented. Our data shows that mice did not retain learned performance, indicating that they relied on the sensory stimuli for expert performance (see Post et al. (2023)). Custom-written routines and Psychtoolbox in MATLAB were used to present the visual stimuli, to trigger the lickport to dispense and retract water, and to acquire data.

### Cranial window surgery

Craniotomies were performed at 6-8 weeks. Prior to surgery, mice were given dexamethasone (0.2 mg/kg) and carprofen (5 mg/kg) intraperitoneally and subcutaneously respectively. Mice were anesthetized with isoflurane (5% induction, 1.5-2% maintenance via nose cone) and placed in a stereotaxic frame. Under sterile conditions, a 4.5 mm diameter craniotomy was drilled over the right primary visual cortex (V1) and covered with a 5 mm glass coverslip (**Fig. 2C**). Before securing the cranial window with a coverslip, we injected 60-100 nl of pGP-AAV-syn-jGCaMP7f-WPRE. A custom U-shaped aluminum bar was attached to the skull with dental cement to head restrain the animal during behavior and calcium imaging. For two days following surgery, mice were given dexamethasone (0.2 mg/kg) daily.

### Viral constructs

pGP-AAV-syn-jGCaMP7f-WPRE were purchased from Addgene and diluted to a working titer of 2e^13^ with 1% filtered Fast Green FCF dye (Fisher Scientific).

### In-vivo two photon calcium imaging

Calcium imaging was performed on a Scientifica 2-photon microscope equipped with a Chameleon Ultra II Ti:sapphire laser (Coherent), resonant scanning mirrors (Cambridge Technologies), a 20X objective (1.05 NA, Olympus), multialkali photmultiplier tubes (R3896, Hamamatsu) and ScanImage software(Pologruto et al., 2003). Stimulus evoked responses of L2/3 neurons in V1 were recorded at 15.2 Hz in 1 field of view. Each field of view (FOV) consisted of a mean of 95.2 pyramidal cells (sd = 38.3). In each animal, imaging was performed at 150-250 μm.

### Data analysis

#### Data analysis for calcium imaging

Calcium-imaging data were analyzed using suite2p (Pachitariu et al., 2017) and custom-written MATLAB routines. All data was then processed using suite2p for image registration, ROI detection, cell labeling, and calcium signal extraction with neuropil correction. Once suite2p had performed a rigid and non-rigid registration and then detected regions-of-interest (ROIs) using a classifier, we manually selected cells using visual inspection of ROIs and fluorescence traces to ensure the cells were healthy. We then used the deconvolved spikes determined by suite2p in our subsequent analysis that used custom-written MATLAB scripts.

#### Movement-related cell removal

Because movement information has increasingly been found in sensory areas, it was important that we remove any artefacts of movement (Zagha et al., 2022), particularly licking-related activity which would not index sensory processing. We thus identified any cells that were associated with lick movements and removed them from our neural data (Post et al., 2023). We additionally performed a locomotion analysis using video of the mice running. We correlated locomtion with neural activity over the trial periods and found no correlations (data not shown).

#### Lick Decoding

A support vector machine (SVM) was used to predict Pref or NP stimuli from licking data. A radial basis function was used as the kernel. The *fitcsvm* function in MATLAB was used. 80% of data was used to train the SVM and 20% to test. Per time bin (.067 s), 1000 machines were generated per mouse which resulted in 1000 accuracy outputs per mouse. Data were then group averaged and plotted with 95% CI. Shuffled data for controls (not shown) was also tested and found to be at chance levels and is available upon request.

#### Network divergence of network state over trials periods

To determine the degree of network divergence across time within each trial outcome, we computed how far apart network states were over time using bootstrapped Euclidean distances between network positions in which each neuron in a given population constituted a dimension, e.g. **Fig 3B**. These values were then normalized by dividing the Euclidean distance by the square root of the total number of dimensions. A random sample of 12 trials was selected and averaged. We decided to use 12 trials as a sample as this was the smallest value that allowed us to reliably generate normal distributions for the bootstrap across mice. 1000 means were computed for each time point per trial outcome per mouse per session. This resulted in each mouse having either a 23x23x1000 in Hit trials (23 time steps due to our sampling rate of 15.2 Hz over 1.4 s) or a 65x65x1000 matrix in CR and FA trials (65 time steps due to our sampling rate of 15.2 Hz over 4.2 s). Matrices were then averaged along the third dimension within mice, then averaged across mice per trial outcome per session.

#### Decoding of network state over trial periods

To determine how similar or different the dynamics of the network was over the trial periods, we used a Multinomial Naive Bayes classifier to determine whether a given time bin’s dynamic was discriminable from another time bin’s, e.g. **Fig. 3C**. We used the *fitcnb* function in MATLAB. We used 80% of data for training and 20% for testing. We performed 1000 iterations for the entire trial period per mouse. This generated a 23x23x1000 matrix of accuracy values in the Pref conditions and a 65x65x1000 matrix in the NP conditions per mouse. Accuracy values were then averaged to generate a 23x23 or 65x65 matrix. Shuffled data for controls (not shown) was also tested and found to be at chance levels and is available upon request.

#### Network divergence between trial outcomes

To determine the degree of network divergence across time within between trial outcomes, we computed how far apart network states were between trial outcomes over the trial periods using bootstrapped Euclidean distances between network positions in which each neuron in a given population constituted a dimension, e.g. **Fig 4A**. These values were then normalized by dividing the Euclidean distance by the square root of the total number of dimensions. A random sample of 12 trials was selected and averaged. We decided to use 12 trials as a sample as this was the smallest value that allowed us to reliably generate normal distributions for the bootstrap across mice. 1000 means were computed for each time point per trial outcome per mouse per session. This resulted in each mouse having either a 23x1000 in Hit trials or a 65x1000 matrix in CR and FA trials. Matrices were then averaged as the grand mean within each session and plotted with 95% CI.

#### Network Decoding and Feature Selection in Trial Outcome Prediction

Multinomial Naive Bayes classifiers were used to predict trial outcome from neural data using the *fitcnb* function in MATLAB. If decoding occurred with a selected group of cells (e.g. **Fig. 6B**, network group), a forward feature selection algorithm was employed to identify a subset of the most informative cells using a 5-fold CV partition. These cells then composed the feature space. Feature selection was employed using the *sequentialfs* function in MATLAB. This was performed 1000 times per cell group per time bin per mouse. If all cells in the network were used, no CV partition was employed, and 1000 decoders were generated per time bin per mouse, e.g. **Fig. 4B**. 80% of the data was used in training and 20% in testing. Averages were then computed using the group mean and plotted with 95% CI. Shuffled data for controls (not shown) was also tested and found to be at chance levels and is available upon request.

#### Correlation of Network Divergence and Decoder

Correlations between decoder accuracy and Euclidean distance in trial outcomes were computed by correlating the mean accuracy and mean Euclidean distance using the Pearson correlation coefficient.

To calculate correlations of network divergence and decoding over trial periods, we used only the values above and below the diagonal, as the Euclidean distance measure yielded a value of 0 in distance between states of the same time. Additionally, because the Euclidean distance values were mirrored along the diagonal, we did not wish to bias the correlations by using Euclidean distance values twice. However, because the Naive Bayes classifier was trained to discriminate network states for every moment of network activity, there were slight differences in accuracy above and below the diagonal. We therefore collapsed each network divergence matrix and each decoder matrix into 2 arrays, one containing values above the diagonal and the other below, and then averaged the arrays. These arrays were then correlated. Matrices in the Pref trial period (e.g. Hit trials) were collapsed from 23x23 matrices to two arrays of 253 values and then averaged. Matrices in the NP trial period (e.g. CR trials) were collapsed from 65x65 matrices to two arrays of 2080 values and then averaged.

Correlations between network divergence between trial outcomes and decoding of trial outcomes was computed using the grand mean of each curve.

#### Single Unit Decoding

Each unit was decoded using Multinomial Naive Bayes classifiers. 80% of data was used to train the classifier and 20% to test. Per cell per time bin (.067 s), 1000 machines were generated per mouse which resulted in 1000 accuracy outputs per cell per mouse. The *fitcnb* function in MATLAB was used.

To plot accuracy curves over the trial period for a given number of cells (e.g. **Fig. 6A**), we selected the most accurate individual cell outputs per time bin per trial per mouse and then averaged them. For instance, at a given time bin in the 4 cell accuracy group, we would take the four cells for a given mouse that are most accurate within a trial and average the accuracies. This leads to 1000 accuracy values per mouse per time bin. These values were then group averaged and plotted with 95% CI.

Shuffled data for controls (not shown) was also tested and found to be at chance levels and is available upon request.

#### Oscillatory Cell Identification

We averaged activity of each cell in Pref and NP trials separately within a session. We then performed a Fast Fourier Transform on the average activity and normalized the spectral density function. If a cell’s spectral density function had only one peak of power above 50%, we considered it an oscillatory cell. Cells identified in this manner were included if they reached this criterion in either or both Pref or NP trials.

#### Ramping Cell Identification

We averaged activity of each cell in Pref and NP trials separately within a session. We then normalized activity and selected cells that had only one peak of activity above 50%. If a cell reached this criterion, it was considered a ramping cell. Cells identified in this manner were included if they reached this criterion in either or both Pref or NP trials.

#### Machine Learning Pipeline

All machine learning was performed using the Nautilus cluster, supported in part by National Science Foundation (NSF) awards CNS-1730158, ACI-1540112, ACI-1541349, OAC-1826967, OAC-2112167, CNS-2100237, CNS-2120019, the University of California Office of the President, and the University of California San Diego’s California Institute for Telecommunications and Information Technology/Qualcomm Institute. Thanks to CENIC for the 100Gbps networks.

### Statistical analyses

All time series data were plotted as the mean with 95% CI. Comparisons between the fraction of oscillatory and ramping cells across sessions were done using Kruskal-Wallis tests, following Lilliefors test of normality.

### Exclusion of mice

In the naive session, 2 mice were excluded from analyses of CR trials as there were few trials (10 and 5). In the middle session, 1 mouse was excluded from all analyses as the only imaging data collected was in naive and learned sessions; 1 other mouse was excluded from analyses of CR trials as there were few trials (8). In the learned session, 1 mouse was excluded from analyses of FA trials as there were few trials (6).

### Data availability

All the analyzed data reported in this study is available from the corresponding author upon request. Additionally, control data for machine learners not shown here is available upon request.

### Code availability

All code used in this manuscript is available from the corresponding author upon request.

## RESULTS

### Changes in V1 neural dynamics accompany learning in the Temporal Pattern Sensory Discrimination Task (TPSD)

We developed a go/no-go task wherein audio-visual patterns were presented to water deprived, awake-behaving mice, as previously described in Post et al. (2023). Stimuli were patterns of 4 synchronous audio-visual stimuli (**Fig. 2A**). The visual stimulus was 90° drifting gratings and the auditory was a 5 kHz tone at 80 dB. Preferred (Pref) and nonpreferred (NP) stimuli differed only in their durations, therefore making TPSD explicitly a temporal discrimination task. A water reward was delivered at 1.2 s from stimulus onset in the Pref condition. The licking window was 1 to 2 s from stimulus onset in both Pref and NP conditions. Mice were placed on a suspended polysterene ball to allow for free movement during the task to reduce stressors and increase performance (**Fig. 2B**) (Guo et al., 2014).

**FIGURE 2:**
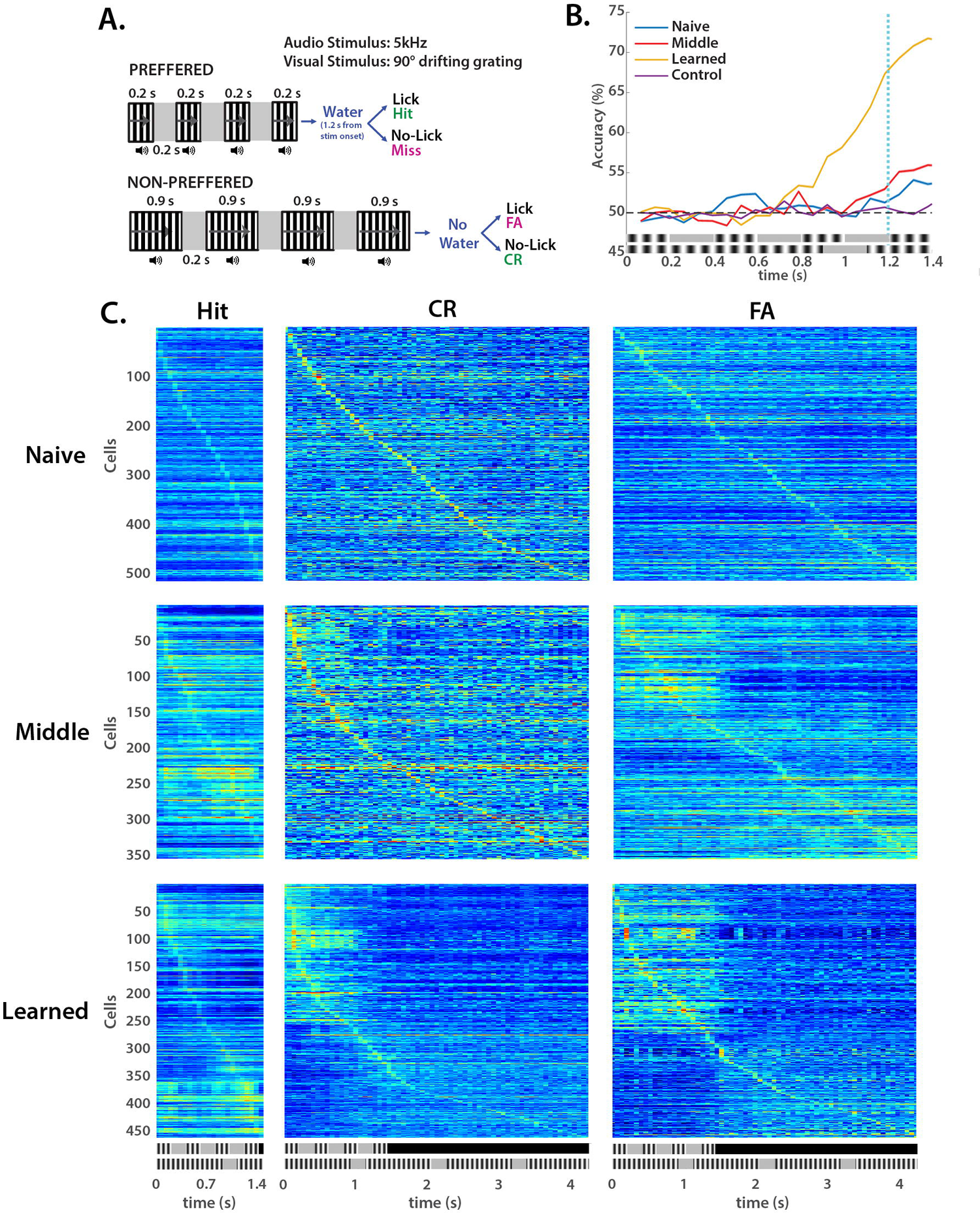
Mice learn Temporal Pattern Sensory Discrimination (TPSD) paradigm and exhibit changes in V1 activity across learning. **A.** Schematic of TPSD. Mice must discriminate subsecond audio-visual patterns based upon their temporal information. **B.** A bootstrapped support vector machine using licking profiles was used to predict Pref or NP stimuli over the trial period to validate learning. Only in the learned session is there sustained difference in licking patterns between conditions prior to the water reward at 1.2 s (blue dotted vertical line). Control sessions are those in which the monitor and speakers are turned off to ensure that mice were not “cheating.” **C.** Spike sorted heatmaps in Hit, CR, and FA trials over sessions show changes in activity dynamics in V1 which suggest circuit restructuring leading to improved performance.

We performed a cranial window surgery in mice in V1 and injected syn-jGCaMP7f (**Fig. 2C**). Upon expression of GCaMP, we had mice perform the TPSD task while simultaneously recording neural activity in V1, L2/3 using 2-photon Ca^2+^ imaging (**Fig. 2D**). As detailed in Post et al. (2023), mice learned the task across sessions, as assessed through a licking decoder (**Fig. 2E**), and exhibited changes in V1 dynamics concomitant with learned performance (**Fig. 2F**). In our previous paper we focused on comparing the neural dynamics between naïve and learned sessions. Here we include neural dynamics changes that occur in middle sessions to assess the role of multiple timing models through the learning process. Using this data, we perform an array of analyses to granularly assess the computational mechanisms of subsecond sensory temporal encoding with respect to prominent theoretical models.

### Network divergence indexes learning and supports the state dependent network model

A prominent model of timing, the state depedent network model (**Fig. 1D**), suggest that temporal information is encoded in the changing population vector of a network over a stimulus’ duration (Mauk & Buonomano, 2004). As sensory information intrinsically carries both spatial and and temporal features, a network’s state will evolve through time based upon its synaptic weights, short-term plasticity, and its intrinsic time constants (Buonomano, 2000; Buonomano & Maass, 2009). This leads to trajectories of the network that are dependent upon a stimulus’ spatial and temporal features.

If temporal information is encoded within the changing population vector of a network as predicted, it follows that the degree of divergence of the population vector would correspond to the degree of behavioral performance in the TPSD task. Divergence would occur within a stimulus, to differentiate one moment in time from another, and between stimuli with different temporal properties. If there is little divergence over time, a stimulus’s temporal properties are not encoded, and if there is little divergence between Pref and NP stimuli, behavioral output would be the same in both conditions. Vice versa, if there is large divergence of the network over the stimulus period, the stimulus’s temporal properties are encoded, and if there is large divergence between Pref and NP stimuli, behavioral output would differ.

To test this prediction, we calculated the Euclidean distance of the network as a measure of network divergence across trial periods (**Fig. 3A,B**). We found that there was an evolution of network divergence across learning in Hit, CR, and FA trials (Miss trials were not included as there were too few samples). In learned sessions, all three trial outcomes displayed increased network divergence from naive sessions. In learned session CR and FA trials, the greatest divergence was seen between the Pref stimulus period (0-1.4 s) and the remaining NP stimulus period, which suggests that mice attended only to the Pref stimulus period regardless of stimulus. However, FA trials exhibited considerably less network divergence than CR trials did in the Pref stimulus period, which may explain why mice responded with a go response in FA trials and withheld in CR trials – that is, temporal information of the stimulus was not accurately encoded in FA trials.

**FIGURE 3:**
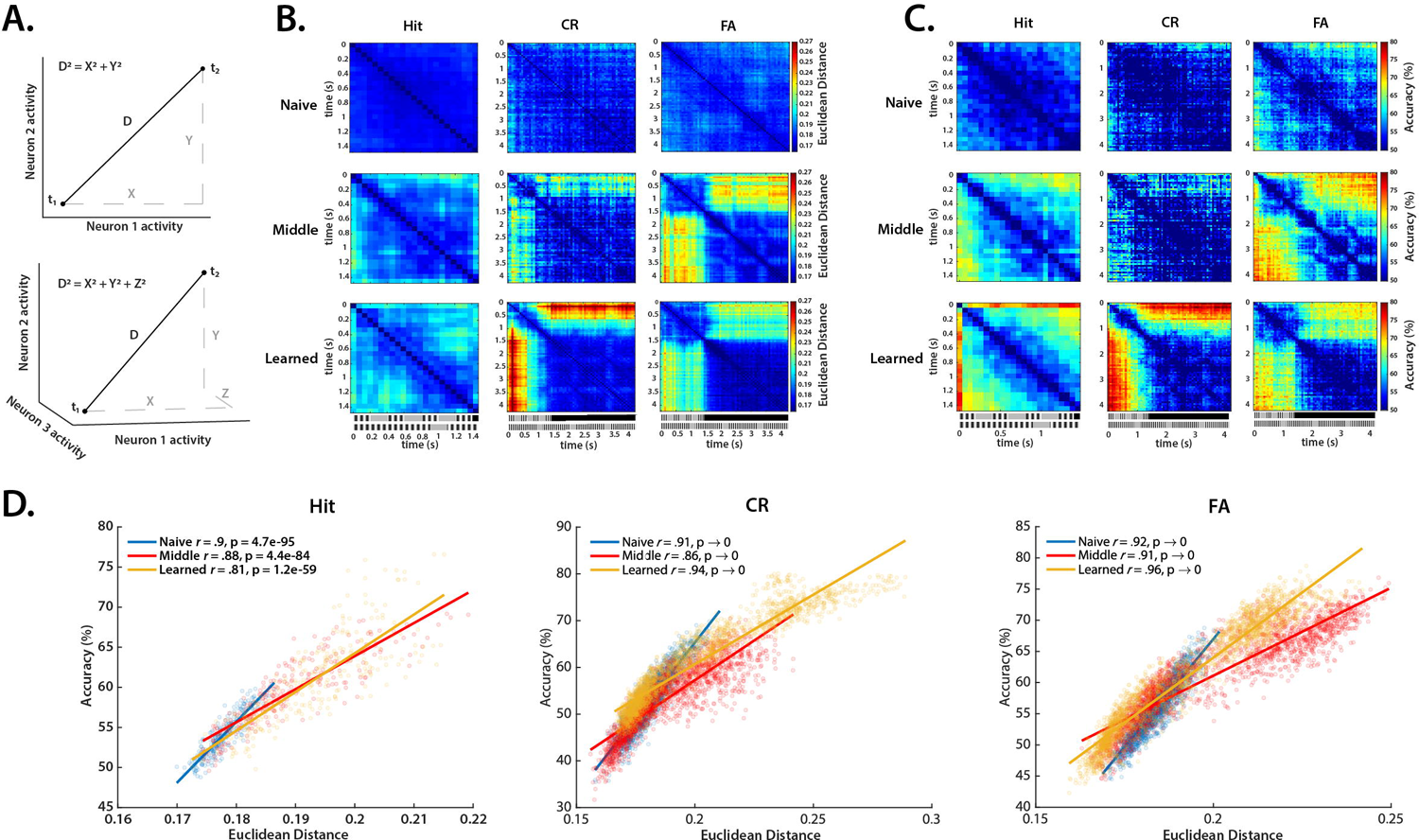
Network divergence and decodability of network state across the trial period increases over learning. **A.** Example of calculation of Euclidean distance. The top panel shows a two neuron system in which the network state changes between *t_1_* and *t_2_*. The distance between the network states (*D*) can be calculated using the Pythagorean theorem, i.e. Euclidean distance in two dimensions. The bottom panel shows the calculation in a 3 dimensional, i.e. 3 neuron system. The equation can be generalized to *n* dimensions (see Methods). **B.** Network divergence across sessions in Hit, CR, and FA trials. Network divergence was calculated as the bootstrapped Euclidean distance between the positions of the network at different points in time (see Methods). **C.** Naive Bayes classifier decoding of network state between different points in time across sessions in Hit, CR, and FA trials. **D.** Correlations of network divergence in A. and network state decoding in B. across sessions in Hit, CR, and FA trials. Pearson’s correlation coefficient was used to calculate correlations.

If network divergence was the computational mechanism of temporal encoding, we predicted that a neural decoder would accurately discriminate network activity over the trial period only when network divergence was high, and that discriminability of network activity would increase with learning. We employed a Naive Bayes classifier to discriminate network activity over trial periods in naive, middle, and learned sessions and found that discriminability of network activity increased with behavioral performance (**Fig. 3C**). The performance of the decoder was highly correlated with network divergence across sessions and trial outcomes (**Fig. 3D**). Notably, the Naive Bayes decoder predicts outcomes through probabilistic learning, a manner of classification distinct from the geometric solutions found through calculation of Euclidean distance. Because each method solves these problems differently but arrives at highly correlated solutions, it is exceedingly likely that state-space trajectories underlie temporal encoding in V1.

We next tested the hypothesis that network divergence between stimuli would predict trial outcome and that the degree of divergence would index behavioral performance. Specifically, we predicted that Hit and CR trials and CR and FA trials would diverge with learning, but that Hit and FA trials would not. Indeed, we found that across learning, Hit and CR trials and CR and FA trials diverged, beginning in the middle sessions and increasing in learned sessions (**Fig. 4A**). Hit and FA trials remained non-divergent throughout naive, middle, and learned sessions (**Fig. 4A**).

**FIGURE 4:**
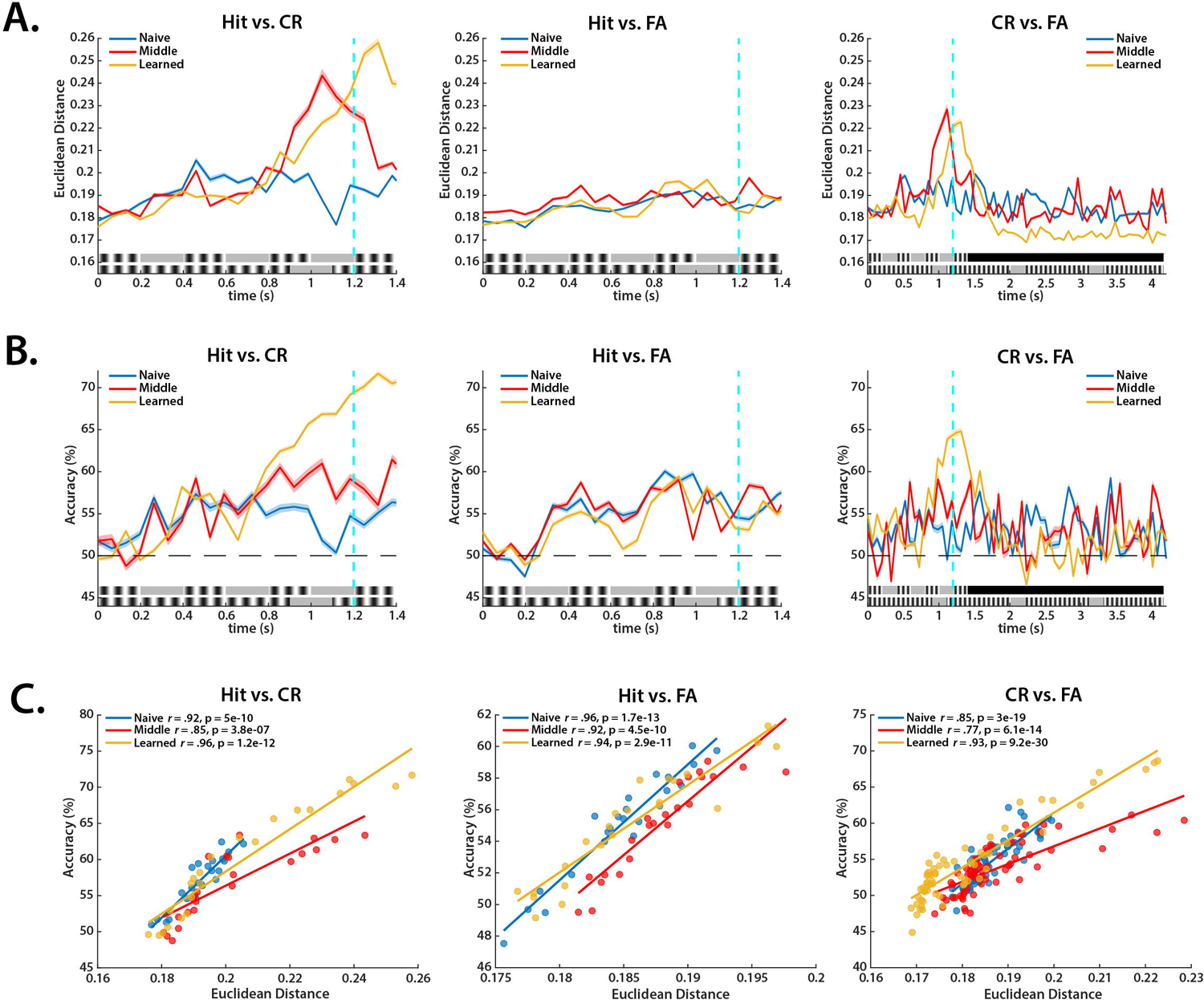
Network divergence and decodability of trial outcomes increases over learning. **A.** Network divergence between trial outcomes in naive, middle, and learned sessions. Network divergence was calculated as the bootstrapped Euclidean distance between trial outcomes at a given time (see Methods). Curves are plotted with 95% CI. **B.** Naive Bayes classifier decoding of trial outcomes over the trial period using neural data. Curves are plotted with 95% CI. **C.** Correlations of network divergence between trial outcomes in A. and trial outcome decoding in B. across sessions. Pearson’s correlation coefficient was used to calculate correlations.

We hypothesized as before that network divergence was correlated with decodability. We used Naive Bayes classifiers to discriminate trial outcomes over the stimulus periods and found that decodability increased with learning and that Hit and CR trials and CR and FA trials were highly discriminable while Hit and FA trials remained at similarly low levels of discriminability across sessions (**Fig. 4B**). We correlated network divergence with the classifiers’ performance and found that across sessions and trial outcomes there were high correlations, evincing again that temporal encoding is achieved through high dimensional neural trajectories (**Fig. 4C**).

### Learning on the TPSD task is supported by encoding temporal information at the level of the network rather than at the single unit level

Although our results provide strong evidence that temporal information was encoded in network trajectories, other mechanisms such as changes in single cell (single unit) activity could account for temporal encoding. Two prominent models of timing, ramping and synfire models, may rely on single unit activity to generate temporal information. Indeed, ramping activity has been found in V1 in reward timing neurons and neurons encoding sensory expectations (Chubykin et al., 2013; Gavornik & Bear, 2014b, 2014a; Monk et al., 2020; Shuler & Bear, 2006).

To test whether our mice relied on single cell activity to encode temporal information in the TPSD task, we used a decoder to predict trial outcomes from neural activity in naive, middle, and learned sessions for each cell and then sorted units by when in the trial period they were most accurate, akin to spike sorted heatmaps (**Fig. 5**). We found that predictability was sparsely tiled across Hit vs. CR, Hit vs. FA, and CR vs. FA trials in naive sessions. Accuracy became less sparse around the water reward period in middle sessions in Hit vs. CR trials and CR vs. FA trials, but not in Hit vs. FA trials. This profile was broadened in learned sessions in which accuracy values were network wide in Hit vs. CR trials and CR vs. FA trials around the water reward and prior to it. Hit vs. FA accuracy values remained sparse across the network in learned sessions. These results suggest that learning recruits the entire network as opposed to encoding time at the single unit level, such as in the manner of synfire chain or ramping models.

**FIGURE 5:**
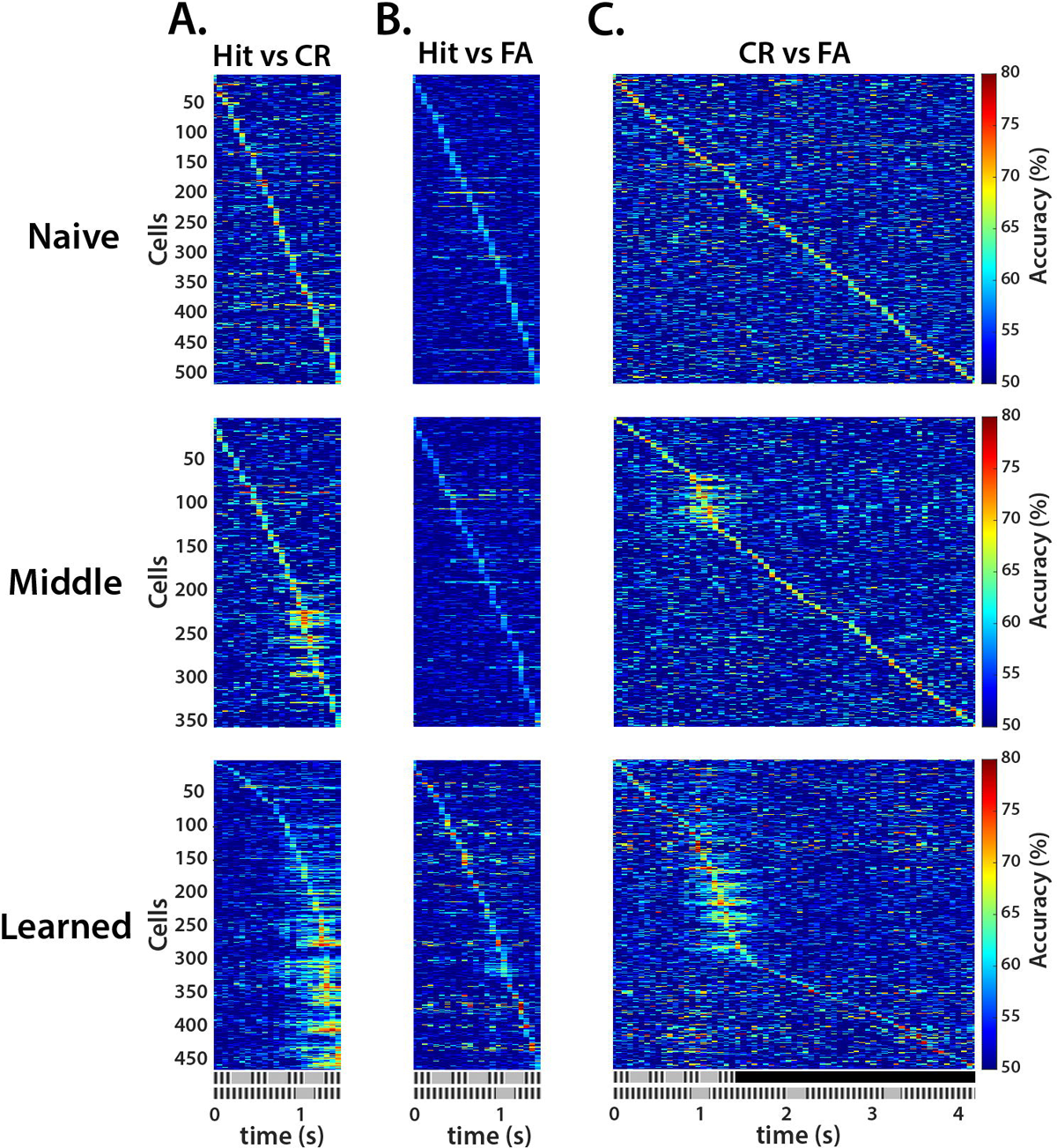
Single unit decoding between trial outcomes. **A.** Each cell was used to discriminate Hit from CR trials across sessions. Cells are then sorted over the trial period by the point at which they were most accurate, akin to spike sorted heatmaps. Naive sessions are the first row, middle sessions the middle row, and learned sessions the bottom row. Bootstrapped Naive Bayes classifiers were used for decoding. **B.** As in A., but Hit vs. FA. **C.** As in A., but CR vs FA.

To compare whether temporal information was better encoded at the network or single unit level, we compared the decoding accuracies of the single unit regime to decoding accuracies of a network regime. To do this we, we first decoded network activity between trial outcomes in which a given number of cells composed the feature space. We iteratively did this in groups of 1 to 80 cells by using a forward feature selection algorithm to find the most informative cells within a network for a given point in time. We then iteratively took the most predictive cells for a given time from the single unit decoding regime in **Fig. 5** and averaged their accuracy scores (see Methods). We then compared network and single unit decodability.

We found that decoding Hit from CR trials (**Fig. 6A**) and CR from FA trials (**Fig. 6C**) increased in accuracy across mice’s learning in the network condition, but decreased in accuracy across sessions in the single unit condition. Decoding Hit from FA trials (**Fig. 6B**) increased in accuracy across sessions in the single unit condition and remained at similar accuracy in the network condition. Because neural decoding accuracy was directly proportional to behavioral performance through learning in the network condition and was inversely proportional to learning in the single unit condition, we conclude that temporal information is encoded at the network level as opposed to the single unit level.

**FIGURE 6:**
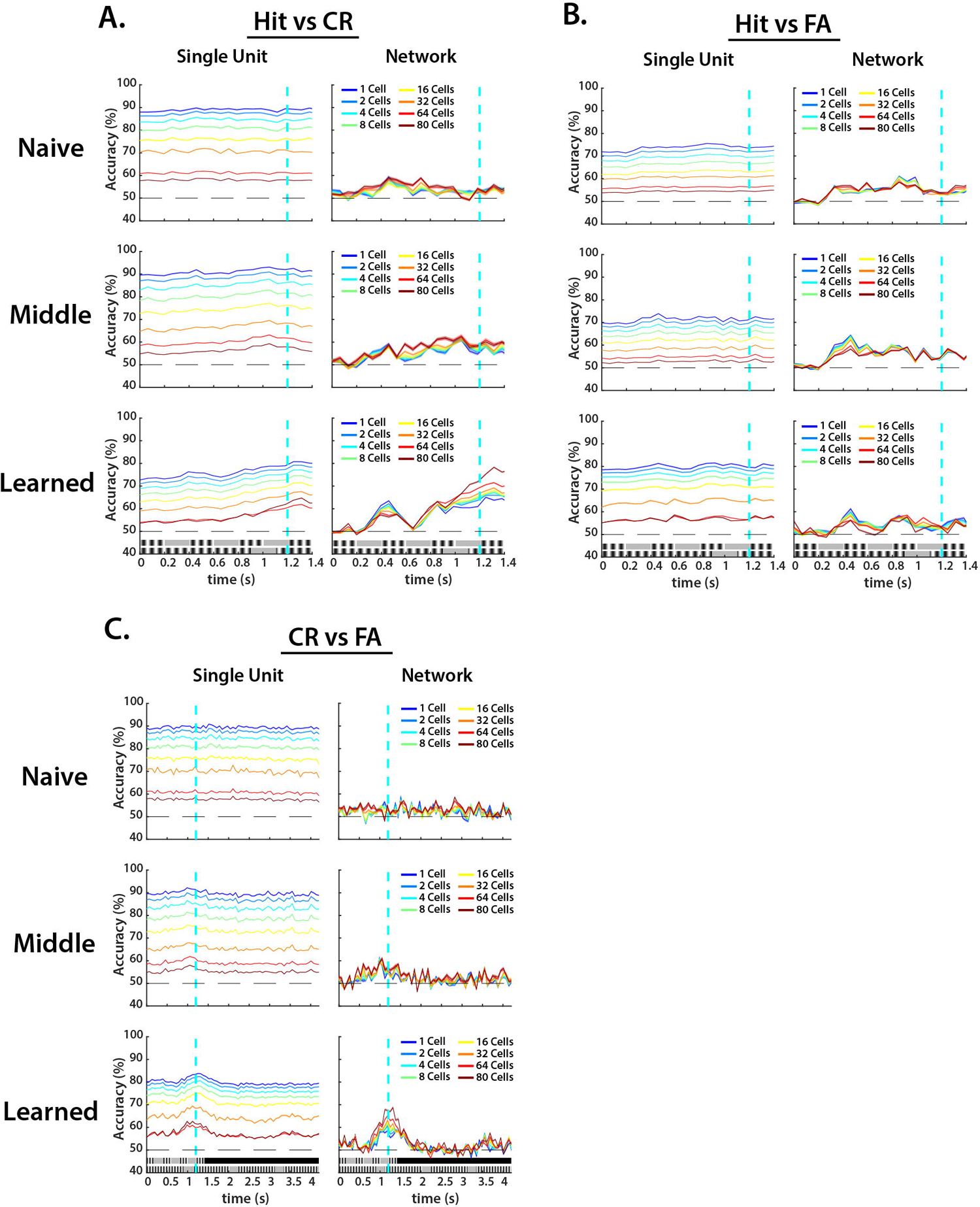
Temporal information is encoded at the population level, not the single unit level. **A.** Decoding Hit from CR trials using neural data as single units from Fig. 5A (left column) or as a network of increasing numbers of cells (right column). Naive sessions are the first row, middle sessions are the middle row, and learned sessions are the bottom row. **B.** As in A., but decoding Hit from FA trials. **C.** As in A., but decoding CR from FA trials. See Methods for details regarding cell selection procedures. All curves are plotted with 95% CI.

### Oscillatory activity does not account for temporal encoding in TPSD task

The first models of timing proposed that oscillatory activity was responsible for temporal encoding. In these models, it was predicted that a pacemaker module, much like a metronome, regularly outputs pulses to a downstream accumulator where they are then counted (Creelman, 1962; Gibbon, 1977; Gibbon et al., 1984). Subsequent models predict that a collection of oscillators output to downstream targets, such as striatum, where aggregated oscillatory activity is decoded (**Fig. 1A**) (Buhusi & Meck, 2005; Matell & Meck, 2004; van Rijn et al., 2014). Although evidence of oscillators has not been found in V1, it is conceivable that they are present and are responsible for temporal encoding and accurate discrimination of Pref and NP stimuli in the TPSD task, particularly as our stimuli were periodic.

To test whether oscillatory models accounted for temporal encoding in V1, we first identified any putative oscillatory cells in our recordings. We did not find a significant difference in the fraction of oscillatory cells identified across sessions (**Fig. 7A**). Nevertheless, we hypothesized that if oscillators were responsible for temporal encoding, oscillatory activity would better predict correct responses, particularly in learned sessions, than non-oscillatory activity would. To test this, we decoded Hit from CR trial outcomes in oscillatory (**Fig. 7B**) and non-oscillatory cell populations in the learned session (**Fig. 7C**). Because there were few oscillatory cells found, we used a feature selection algorithm to iteratively select groups of cells that were most informative in both populations so as to avoid any bias in differences in dimensionality. We found comparable levels of decodability in oscillatory and non-oscillatory populations, suggesting that temporal information is not exclusively encoded by oscillatory activity, but is in fact network wide.

**FIGURE 7:**
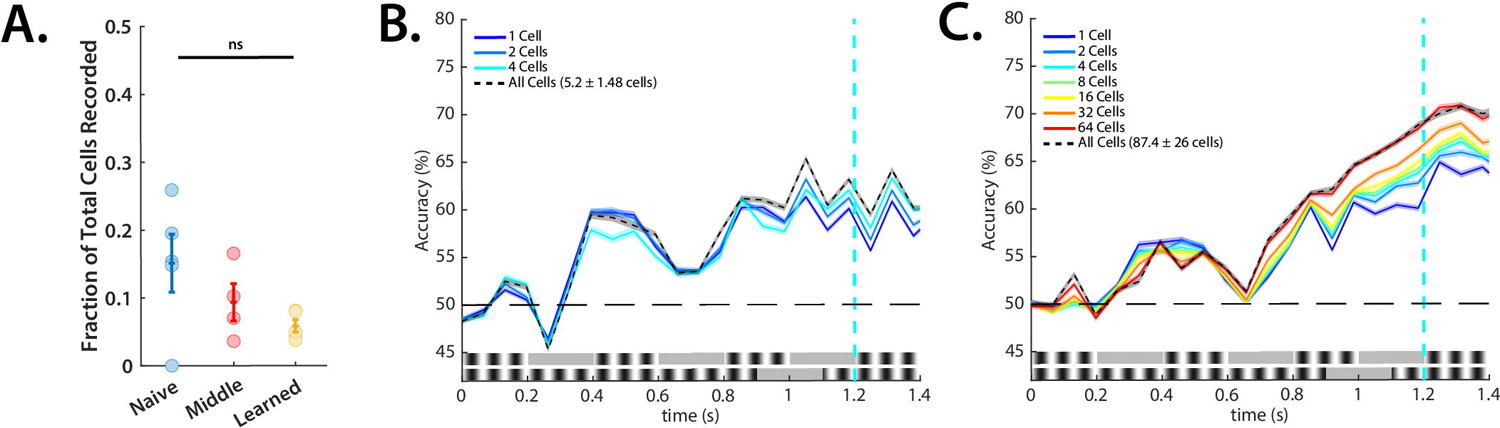
Oscillatory and non-oscillatory activity encode temporal information equally well. **A.** Fractions of oscillatory cells did not significantly change across learning (Kruskal-Wallis, *H*(13) = 2.79, *p* = .25). **B.** Naive Bayes classifier decoding of oscillatory cells in Hit vs. CR trials in the learned session. A feature selection algorithm was used to iteratively select the most informative cells in the population (see Methods). The total network decodability is also shown in the dashed black line. Curves are plotted with 95% CI. **C.** Naive Bayes classifier decoding of non-oscillatory cells in Hit vs. CR trials in the learned session. A feature selection algorithm was used to iteratively select the most informative cells in the population. The total network decodability is also shown in the dashed black line. Curves are plotted with 95% CI.

We therefore hypothesized that oscillatory activity intrinsically emerged from network activity and was an aspect of the network’s state space trajectory. To determine whether this was the case, we calculated the network divergence of the oscillatory population over the trial period (**Fig. 8A**) and then decoded the oscillatory population activity over the trial period (**Fig. 8B**), and then correlated these values (**Fig. 8C**).We additionally calculated network divergence of the oscillatory population between trial outcomes (**Fig. 9A**) and then decoded neural activity between trial outcomes (**Fig. 9B**) and then correlated these values (**Fig. 9C**). We indeed found high correlations across sessions and trial outcomes, signifying that the oscillatory population was in fact operating as a part of the network trajectory in high dimensional state space.

**FIGURE 8:**
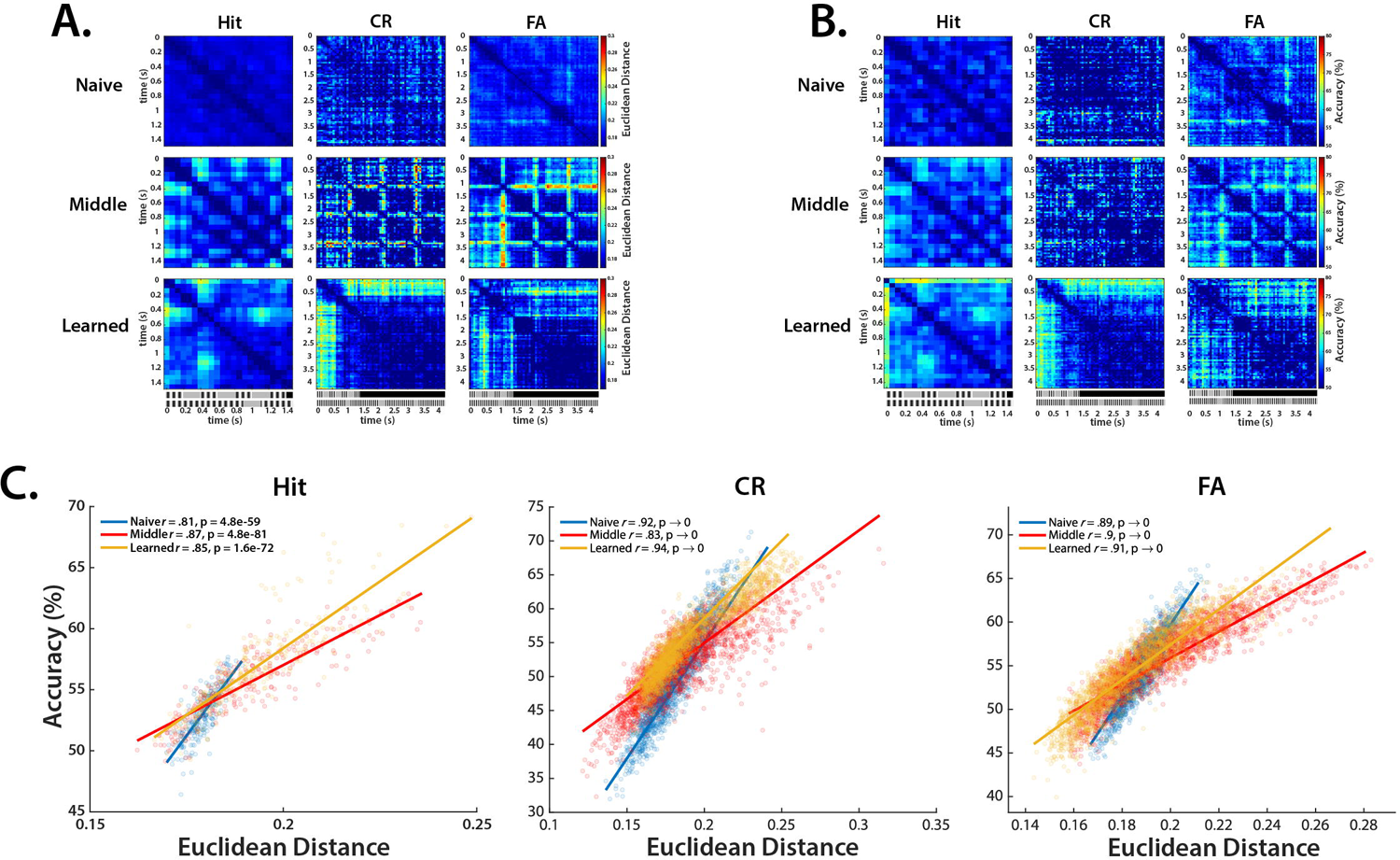
Evolution of oscillatory activity through the trial period is an aspect of network divergence. **A.** Network divergence across sessions in Hit, CR, and FA trials in oscillatory populations. Network divergence was calculated as the bootstrapped Euclidean distance between the positions of the network at different points in time (see Methods). **B.** Naive Bayes classifier decoding of network state between different points in time across sessions in Hit, CR, and FA trials in oscillatory populations. **C.** Correlations of network divergence across the trial period from Fig. 8A and network state decoding from Fig. 8B across sessions in Hit, CR, and FA trials in oscillatory populations. Pearson’s correlation coefficient was used to calculate correlations.

**FIGURE 9:**
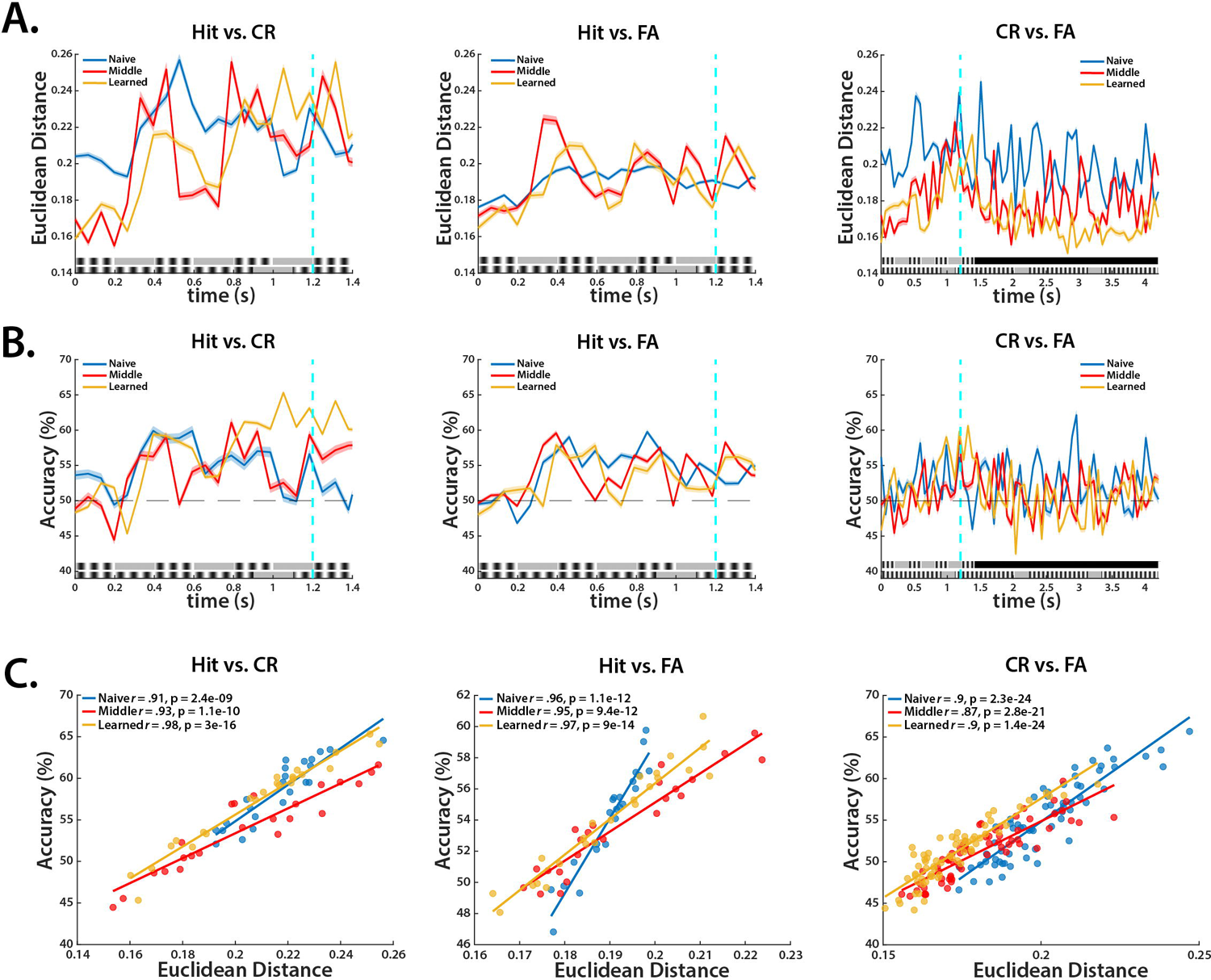
Oscillatory activity differs between trial outcomes according to degree of network divergence. **A.** Network divergence between trial outcomes in naive, middle, and learned sessions in oscillatory populations. Network divergence was calculated as the bootstrapped Euclidean distance between trial outcomes at a given time (see Methods). **B.** Naive Bayes classifier decoding of trial outcomes over the trial period using neural data in oscillatory populations. Curves are plotted with 95% CI. **C.** Correlations of network divergence between trial outcomes from Fig. 9A and trial outcome decoding **and** Fig. 9B across sessions in oscillatory populations. Pearson’s correlation coefficient was used to calculate correlations.

### Ramping activity does not account for temporal encoding in TPSD task

As with oscillatory cells, we sought to test whether ramping activity could account for temporal encoding in the TPSD task (**Fig. 1B**). Unlike oscillators, ramping activity has been found in V1 previously (Chubykin et al., 2013; Monk et al., 2020; Namboodiri et al., 2015; Shuler & Bear, 2006) so it may have been the case that ramping activity better accounted for temporal learning than did network divergence.

We first identified any cells with ramping-like activity and found that across learning there was not a significant difference in the fraction of ramping cells in a given population (**Fig. 10A**). To test whether ramping cells exhibited more informative temporal activity, we used a decoder with a feature selection algorithm to discriminate Hit from CR trials in learned sessions in ramping and non-ramping populations. We predicted that if temporal information was encoded in ramping activity, decodability in the ramping population would be greater than in the non-ramping population. We iteratively selected groups of cells from each population over the trial period to avoid biases of greater dimensionality in the non-ramping population.

**FIGURE 10:**
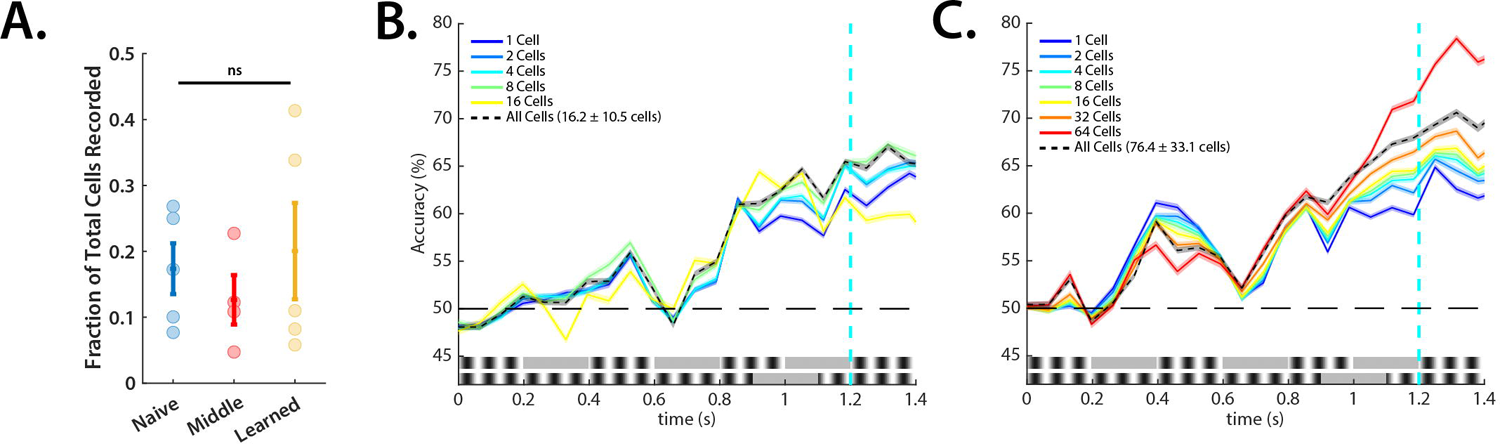
Ramping and non-ramping activity encode temporal information equally well. **A.** Fractions of ramping cells did not significantly change across learning (Kruskal-Wallis, *H*(13) = .5, *p* = .78). **B.** Naive Bayes classifier decoding of ramping cells in Hit vs. CR trials in the learned session. A feature selection algorithm was used to iteratively select the most informative cells in the population (see Methods). The total network decodability is also shown in the dashed black line. Curves are plotted with 95% CI. **C.** Naive Bayes classifier decoding of non-ramping cells in Hit vs. CR trials in the learned session. A feature selection algorithm was used to iteratively select the most informative cells in the population. The total network decodability is also shown in the dashed black line. Curves are plotted with 95% CI.

We found that ramping and non-ramping populations encoded temporal information comparably (**Fig. 10B-C**). We therefore hypothesized that, as with oscillatory activity, ramping activity intrinsically emerged from network activity and was an aspect of the network’s state space trajectory. To determine whether this was the case, we calculated the network divergence of the ramping population over the trial period (**Fig. 11A**) and decoded the ramping population activity over the trial period (**Fig. 11B**), and then correlated these values (**Fig. 11C**). We additionally calculated network divergence of the ramping population between trial outcomes (**Fig. 12A**) and then decoded neural activity between trial outcomes (**Fig. 12B**) and correlated these values (**Fig. 12C**). We consistently found high correlations across sessions and trial outcomes, signifying that the ramping population was likely was an aspect of the network trajectory through high dimensional state space.

**FIGURE 11:**
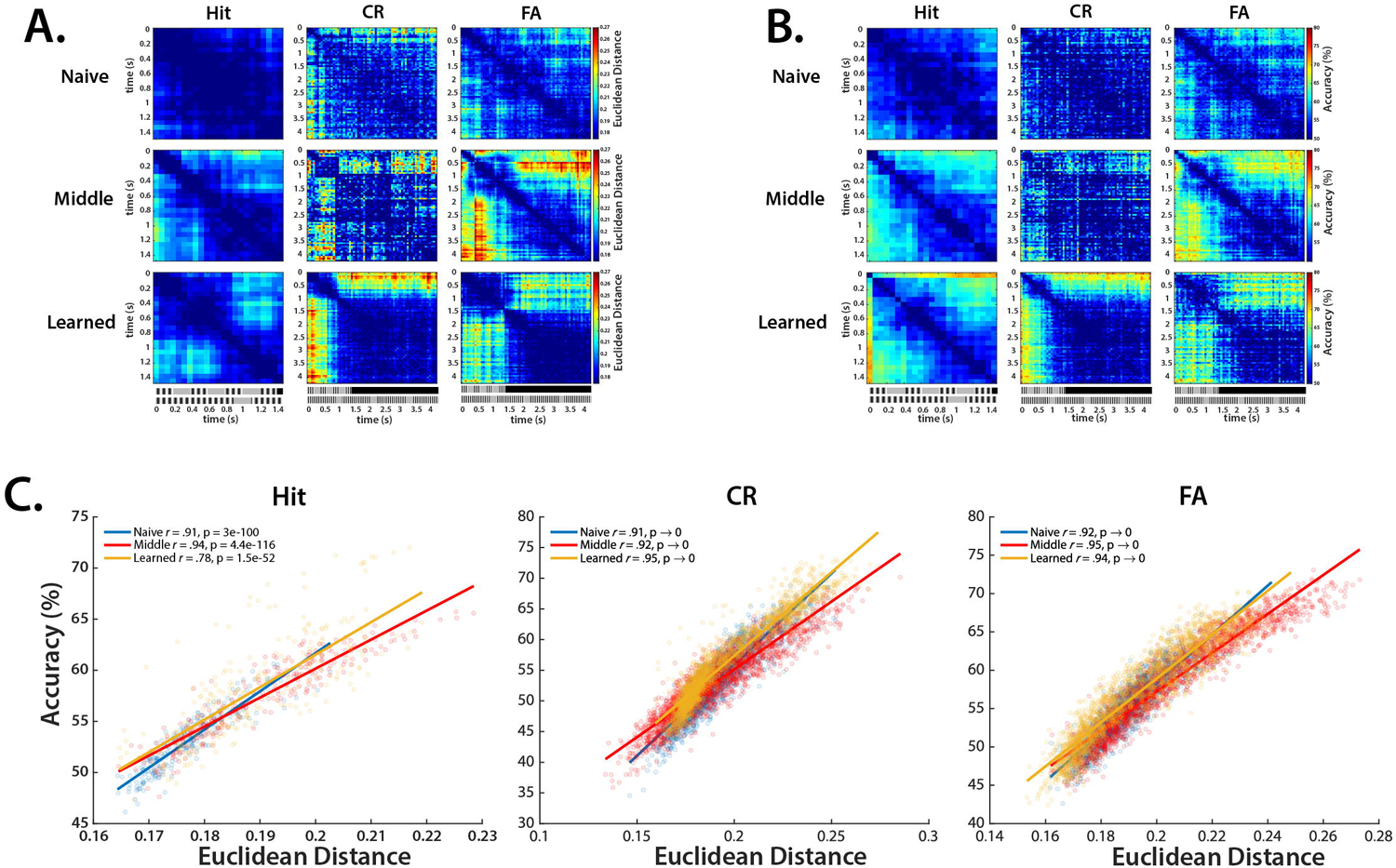
Evolution of ramping activity through the trial period is an aspect of network divergence. **A.** Network divergence across sessions in Hit, CR, and FA trials in ramping populations. Network divergence was calculated as the bootstrapped Euclidean distance between the positions of the network at different points in time (see Methods). **B.** Naive Bayes classifier decoding of network state between different points in time across sessions in Hit, CR, and FA trials in ramping populations. **C.** Correlations of network divergence across the trial period from Fig. 11A and network state decoding from Fig. 11B across sessions in Hit, CR, and FA trials in ramping populations. Pearson’s correlation coefficient was used to calculate correlations.

**FIGURE 12:**
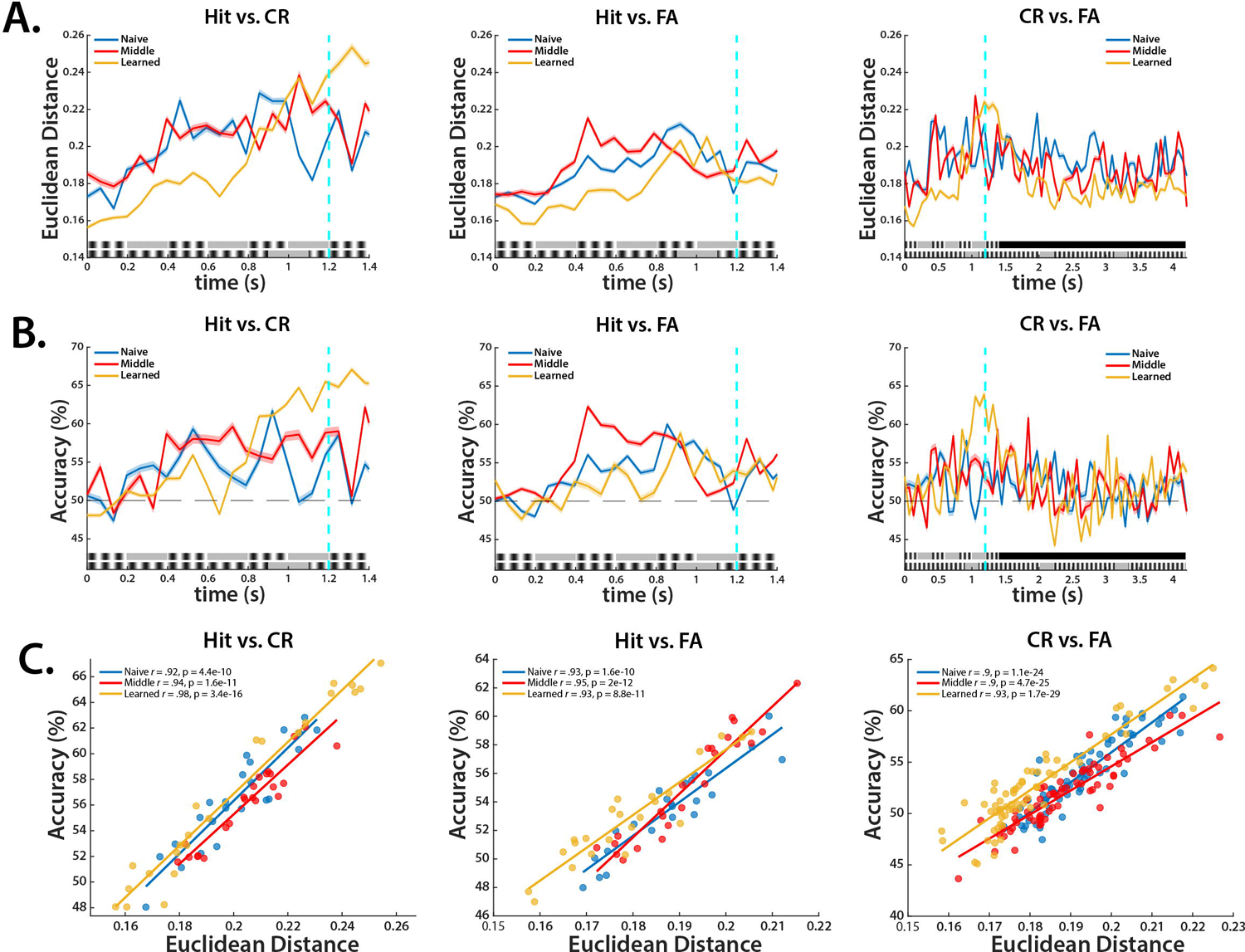
Ramping activity differs between trial outcomes according to degree of network divergence. **A.** Network divergence between trial outcomes in naive, middle, and learned sessions in ramping populations. Network divergence was calculated as the bootstrapped Euclidean distance between trial outcomes at a given time (see Methods). **B.** Naive Bayes classifier decoding of trial outcomes over the trial period using neural data in ramping populations. Curves are plotted with 95% CI. **C.** Correlations of network divergence between trial outcomes from Fig. 12A and trial outcome decoding from Fig. 12B across sessions in ramping populations. Pearson’s correlation coefficient was used to calculate correlations.

## DISCUSSION

Though time as a dimension in stimulus encoding has been largely overlooked, it is an integral component. The notes of a song, for instance, can be played in perfect sequence, but if the temporal structure between them is aberrant and chaotic, the song loses its identity. Prey-predator interactions perhaps best capture how critical temporal perception is: not only must a prey or predator anticipate where its counterpart will be, but *when*. The lion is not successful if it occupies the location of its prey from 200 ms ago, or 1 second from where the prey will be in the future. The predator must occupy the same space at the same time as its prey, and this process necessarily entails encoding stimuli from the present and the immediate past in order to anticipate events of the future.

Several models of timing at this scale – milliseconds to seconds – have been proposed that largely can be construed as either dedicated or intrinsic models. We sought to test which of these models best captures neural activity in V1 in mice performing a temporal discrimination task in which audiovisual stimuli differed only in their temporal information. We found considerable evidence that temporal information in the millisecond range is encoded by high dimensional neural trajectories. We examined neural data across sessions and found that in learned sessions, the network’s activity was far more divergent than in naive sessions. Further, between correct trial outcomes, we found that this divergence was maximized. This network divergence was highly correlated with a number of decoding schemes we used, which suggests that the decoders independently recognized and exploited network divergence as an informative coding schema. Even among other proposed models of timing, namely oscillatory and ramping models, we found that network divergence was highly correlated with decodability, implying that these types of activity were in fact aspects of the network divergence of the entire network as opposed to specialized, dedicated mechanisms of timing.

Although V1 has historically been understood as extracting low-level spatial features from visual information, recent evidence has suggested it processes temporal information as well. Shuler and Bear (2006) found evidence of reward timing in V1. Gavornik and Bear (2014b) later found that V1 encodes sequences of stimuli in a temporally-defined, predictive manner. Spatiotemporal prediction has also been found in V1 in mice performing foraging tasks in virtual reality (Fiser et al., 2016; Yu et al., 2022). Nevertheless, these findings did not explicitly test temporal processing in a sensory discrimination task, and an outstanding question was how temporal information was computationally encoded in V1. We tested this and found compelling evidence that temporal processing in V1 follows the state dependent network model in which temporal encoding occurs through the evolution of a network’s population vector in state space. Notably, no one specialized group of cells contained greater temporal information than non-specialized cells.

We found evidence of ramping activity, which accords with previous findings of reward timing in V1. However, ramping activity associated with reward prediction is cholinergically mediated, which may induce state changes in V1 but not changes in temporal processing of stimuli per se (Chubykin et al., 2013; Shuler & Bear, 2006). Furthermore, we removed any cells from our population that were associated with licking that may, as an artefact, have exhibited ramping like activity. The remaining cells that exhibited ramping activity in our recordings likely were recruited by the network as the trial period progressed in order to support the divergence of network states. It has been found that orientation-selective cells in V1 can shift their tuning curves through leaning, and as only ∼40% of V1 cells are simple cells (Cossell et al., 2015; Froudarakis et al., 2019; Kondo et al., 2016), the remaining population may have been preferentially recruited to support learning and push the network to different attractor basins.

We also found evidence of oscillatory activity, although in learned sessions, there were only a handful of oscillatory cells. This was surprising as our stimuli were periodic. One may suspect that activity in orientation selective cells tuned to our gratings would activate in a periodic fashion, and indeed, average activity of the network supports this hypothesis (Post et al., 2023). However, at the single unit level, this was not found to be the case. Instead, our results suggest that temporal information was encoded through the evolution of population activity in both oscillatory and non-oscillatory populations. Our results do not rule out the possibility of a centralized oscillator however. It may be the case that V1 is reading out the activity of an upstream oscillator as high dimensional trajectories. This would require the oscillator to receive visual information from non-cortical areas and then project temporal information to V1 to be reintegrated with spatial information. Biologically, this seems an unlikely mechanism however.

It has been shown that organotypic cortical slices are capable of “learning” a duration, which suggests that intrinsic mechanisms can support subsecond temporal encoding (Goel & Buonomano, 2016). In the slices, polysynaptic activity increased in a temporally dependent manner, and inhibition was suppressed at the learned duration. This suggests that a complex interplay of recurrent excitation and feedforward inhibition can generate population activity that evolves over a trained period to represent elapsed time. In fact, it has been proposed that the differing dynamics of temporal encoding in cortex and striatum are attributable to their connectivity motifs – recurrent excitation in cortex leads to high dimensional trajectories, and recurrent inhibition in striatum leads to sparse, winner-take-all sequentiality (Bakhurin et al., 2017). This may be why we did not find evidence for sparse temporal encoding in V1 as predicted by the synfire chain model.

Indeed, inhibition has been found to be critical in encoding temporal information in a recurrent neural network model (Zhou et al., 2022). However, inhibitory activity is considerably diverse, with GABAergic cells differing in firing profiles, baseline excitability, morphologies, and preferences in where to synapse. In the cortex, parvalbumin (PV), somatostatin (SST), and vasointestinal peptide (VIP) cells are the primary inhibitory interneuron subtypes (Cardin, 2018; Kullander & Topolnik, 2021), and their functional diversity can broaden the encoding space of a network. In a model of a cortical microcircuit, adjusting the synaptic weights of PV and SST inhibitory interneurons onto Pyramidal (Pyr) cells generated an array of Pyr firing profiles in a temporally defined manner, which was attributed to differing short-term plasticity profiles of each cell type (Seay et al., 2020). Experimentally, SST activity has been found in motor cortex to structure sequential activity in a learned motor task (Adler et al., 2019). Similar to Goel and Buonomano (2016), it was found that inhibitory activity viz. SST cells was suppressed through learning and then returned to baseline following learning to structure network activity. Because SST cells in L2/3 of cortex synapse preferentially with Pyr dendrites in L1 (Urban-Ciecko & Barth, 2016; Wu et al., 2023), SST cells through fine, dendritic computation may orchestrate ensembles of Pyr activity in a temporally defined manner which leads to emergent network trajectories over time.

Population vector encoding is exploited as a computational strategy across the brain, possibly due to the increased informational space available. However, in behaviors typically associated with population encoding such as olfaction (Canto-Bustos et al., 2022; Oswald & Urban, 2012), motor output (Georgopoulos et al., 1986; Georgopoulos & Carpenter, 2015), and memory (Grewe et al., 2017; Lee et al., 2023), temporality is implicit, and it is unclear if a network can induce coherent population codes in a time depedent manner as the state dependent network model proposes. In memory in particular, temporality is integral, and it remains to be determined if the simultaneous activation of an ensemble or if sequential activation of a set of ensembles encodes the content of a memory and its duration. Our results suggest that temporal information can emerge through the sequential activation of ensembles, such that the network state diverges across time. Notably, this network state divergence emerges through learning and reliably indexes trial outcome throughout sessions.

Our findings add to a growing body of literature that supports the state dependent network model of timing and finds that temporal information can be encoded intrinsically and mediated by a circuit’s local parameters. Our results provide further evidence that temporal information is encoded by the brain in lower order areas and suggest that time is an integral component of sensory processing.

## ACKNOWLEDGEMENTS

We thank Bart Kats for help with using the Nautilus clusters for running the machine learners.

